# WDR62 and CEP170 recruit MAPKBP1 for pericentriolar material cohesion and mitotic spindle formation

**DOI:** 10.64898/2026.02.23.707558

**Authors:** Uda Y. Ho, Yvonne Y. Yeap, Cheng-Hui Goh, Peter G. Noakes, Dominic C.H. Ng

**Author notes:** Corresponding authors: Uda Ho and Dominic Ng.

## Abstract

Centrosomal and microtubule-associated proteins such as CEP170 and WDR62 are essential in regulating mitotic spindle formation and pole orientation during cell division. MAPKBP1, a paralog of WDR62, is also a centrosomal protein, but its function is currently unclear. We have shown here that MAPKBP1 is localised to the subdistal appendages of the mother centriole, the pericentriolar material (PCM) of the centrosomes and the mitotic spindles during metaphase. Furthermore, MAPKBP1, WDR62 and CEP170 exists as a complex, where MAPKBP1 is recruited to the centrosomes by WDR62 and CEP170, and CEP170-MAPKBP1 interaction is mediated by WDR62. In addition, MAPKBP1 depletion leads to mitotic spindle defects and delayed mitosis that were further exacerbated with WDR62 knockout, indicating a possible redundancy between MAPKBP1 and WDR62. MAPKBP1 loss also leads to PCM fragmentation, which supports its role as a subdistal appendages protein vital in maintaining centrosome structure and PCM cohesion for proper anchoring of mitotic spindles. This study provides insight into how subdistal appendages and centrosome and microtubule associated proteins co-operate to tightly regulate mitotic spindle formation and stability.

## INTRODUCTION

Proper formation of mitotic spindles, poles, and their orientation are essential for synchronous chromosome segregation and cell cycle progression. Mitotic spindle poles consist of the centrosome which forms the microtubule organising centre (MTOC). It is comprised of a mother and daughter centriole joined by interconnecting fibres and surrounded by the pericentriolar material (PCM). The mother centriole consists of distal and subdistal appendages that anchor the minus ends of the microtubules (Gonczy, 2015; Nigg & Raff, 2009). The mother centriole also forms the base of the primary cilium, known as the basal body, at G0 phase of cell cycle (Breslow & Holland, 2019). The centrioles duplicate alongside DNA during S-phase. Centriole under duplication has been reported in congenital diseases such as microcephaly and ciliopathies, whereas centriole amplification is frequently seen in cancers (Phan & Holland, 2021; Purkerson et al., 2024).

There are many microtubules and centrosome-associated proteins that regulate mitotic spindle formation and pole orientation, including WDR62 and CEP170. WDR62 (also known as MCPH2) is a 166kDa scaffold protein that localises to the proximal centriole during S phase and spindle poles during M phase (Jayaraman et al., 2016; Kodani et al., 2015; Lim et al., 2016; Lim et al., 2015). At G0 phase, it is localised to the basal body of the primary cilium (Shohayeb et al., 2019). Its N-terminus contains WD40 repeats that form a barrel structure for microtubule binding (Huang et al., 2021). By contrast, its C-terminus is phosphorylated by Aurora Kinase A (AURKA) and JNK to regulate WDR62 association or disassociation with the spindle respectively (Lim et al., 2015). WDR62 has been shown to recruit CPAP and IFT88 for primary cilia assembly (Shohayeb et al., 2019), WDR62-CEP170-KIF2A interactions for primary cilium disassembly (Zhang et al., 2019) and WDR62-Katanin interaction for microtubule severing during chromosome segregation (Guerreiro et al., 2021; Huang et al., 2021). Mutations in WDR62 have been reported in primary microcephaly patients (Farag et al., 2013; Nardello et al., 2018; Nicholas et al., 2010; Yu et al., 2010). It has been proposed that WDR62 maintains the pool of radial glial cells during brain development, as WDR62 loss leads to premature progenitor cell differentiation and thinning of the cortex (Bilguvar et al., 2010; Chen et al., 2014; Nicholas et al., 2010; Yu et al., 2010). WDR62 is also involved in oocyte, spermatid, heart and skeletal muscle development (Hao et al., 2022; Ho et al., 2021; Zhou et al., 2018).

WDR62 can homodimerise and heterodimerise with its paralog, mitogen activated protein kinase binding protein 1 (MAPKBP1, also known as JNK binding protein 1) to enhance JNK signalling (Cohen-Katsenelson et al., 2011; Macia et al., 2017; Schonauer et al., 2020; Wasserman et al., 2010). MAPKBP1 is a 168 kDa scaffold protein that shares many structural similarities with WDR62, including the N-terminal WD40 repeat domain, C-terminal JNK binding domain and coiled-coil containing region for heterodimerisation (Macia et al., 2017). Like WDR62, MAPKBP1 also localises to the basal body of primary cilium at resting phase and spindle poles during mitosis (Macia et al., 2017; Schonauer et al., 2020). MAPKBP1 mutations have been reported in nephronophthisis patients (Al-Hamed et al., 2021; Macia et al., 2017; Schonauer et al., 2020), which is classified as ciliopathy. However, the loss of MAPKBP1 function do not overtly affect primary cilium formation (Macia et al., 2017). A recent study shows that JNK regulates MAPKBP1 localisation, and the loss of MAPKBP1 results in cilia disassembly by removing the scaffold that restrains JNK signalling at the basal body (Findeisen et al., 2025). It is still unclear if MAPKBP1 and WDR62 share redundant functions.

CEP170 is a 170kDa protein that contains the N-terminal forkhead associated domain for protein-protein interaction, and C-terminal phosphorylation sites responsible for centrosome and microtubule localisation (Guarguaglini et al., 2005). CEP170 is localised to the subdistal appendages of the mother centriole where it anchors the minus ends of microtubules and promotes microtubule organisation (Kodani et al., 2015). It is also localised to the proximal centrioles where it interacts with C-NAP1 and Ninein for centriole cohesion (Chong et al., 2020). CEP170 can interact with dynein2 to support intraflagellar transport during primary cilium assembly (Weijman et al., 2024). It has been reported that CEP170-KIF2B association and WDR62-CEP170-KIF2A interactions lead to mitotic spindle and primary cilium disassembly respectively (Welburn & Cheeseman, 2012; Zhang et al., 2019). Depletion of CEP170 has been shown to cause spindle misorientation and impaired astral microtubules stability (Pillai et al., 2015; Ren et al., 2025). Furthermore, WDR62-CEP170 interaction is required to maintain meiotic spindle stability in the spermatocytes (Qin et al., 2019). However, how CEP170, WDR62 and/or MAPKBP1 are involved in co-regulating mitotic spindle formation, stability and orientation remain to be elucidated.

This study aims to characterise MAPKBP1 and its interaction with WDR62 and CEP170 in regulating mitotic spindle formation during mitosis. We have shown that MAPKBP1 is a subdistal appendages protein downstream of CEP170 and WDR62. MAPKBP1 loss leads to mitotic spindle defects and delayed mitosis, that is worsened with WDR62 knockout (**KO**), indicating a possible redundancy between these proteins. MAPKBP1 depletion also resulted in PCM fragmentation, whereas WDR62 KO cells displayed both PCM fragmentation and centrosome mis segregation during mitosis suggesting that these paralogue proteins may diverge in functions. Therefore, WDR62, CEP170 and MAPKBP1 form a complex to tightly regulate PCM integrity and microtubules nucleation at the centrosomes for proper mitotic spindle formation.

## METHODS

### Cell culture, transfections, nocodozole and stress treatments

HeLa and AD293 cells were cultured in high glucose DMEM medium (Gibco) supplemented with 10% FBS (Hyclone) and 1% penicillin-streptomycin (Gibco). WDR62 KO AD293 cells were generated using CRISPR technology as previously described (Lim et al., 2016). 2 × 10^5^ cells were seeded to each well in the 6 well plate with or without coverslips. The next day, siRNA or plasmid transfection was performed using lipofectamine 2000 according to manufacturer’s instructions (Invitrogen). The WDR62 WT and mutant plasmids, siRNA sequences and final concentration used are listed in Supplementary Table 1. Microtubules were depolymerised with 10 μM nocodazole treatment for 30 mins at 4°C. For oxidative stress treatment, 1mM H_2_O_2_ (Chemsupply) were added into the media for 30 mins or 60 mins at 37°C. For osmotic stress treatment, 0.5M sorbitol (Sigma) were added into the media for 30 mins or 60 mins at 37°C. 50μM JNK inhibitor VIII (Calbiochem) were added into the media for 30 mins or 60 mins at 37°C.

### Immunofluorescence

Immunofluorescence was performed as previously described (Lim et al., 2015). Cells were fixed in 100% ice cold methanol. Cells were permeabilised with 0.2% Triton-X for 20 mins and blocked in blocking buffer (20% FBS, 0.2% Triton-X in phosphate buffered saline pH 7.4 (**PBS**) for 30 mins before incubating with primary antibody at 4°C for 1 h at room temperature. After three PBS washes, cells were incubated with appropriate secondary antibodies and counterstained with DAPI before mounting in Prolong Diamond (Invitrogen). Negative controls (using IgG antibody or no primary antibody) were included in each experiment. The primary and secondary antibodies, IgG control antibodies used, and their dilutions are listed in Supplementary Tables 2 and 3. Microscopy was performed using Leica DMi8 SP8 inverted confocal microscope and Z-stacks were collected at 0.5μm intervals. Z-stacked maximum intensity projection images were compiled using the Leica Application Suite Advanced Fluorescence Lite (LAS AF Lite) software version 2.6.3 build 8173 (Leica Microsystems).

### Immunoblotting

Cells were lysed in RIPA buffer (50mM Tris-HCl pH7.3, 150mM NaCl, 0.1mM EDTA, 1% sodium deoxycholate, 1% Triton X-100, 0.2% NaF and 100μM Na_3_VO_4_ supplemented with complete protease inhibitors (Roche) and quantitated using Bradford assay (BioRad) against BSA standard. Immunoblotting was performed as previously described (Lim et al., 2016). Briefly, 50μg of protein lysates were ran on the 4-15% Tris-glycine gradient precast gel (BioRad), transferred to PVDF membrane (Millipore) and blocked in 5% skim milk, before incubating with primary antibody at 4°C overnight. The next day, membrane was incubated with secondary antibody for 1 h and bands were detected using ECL Clarity (BioRad), visualised using Odyssey-Fc and bands were quantified using Image Studio Lite version 5.2. The primary and secondary antibodies used, and their dilutions are listed in Supplementary Tables 2 and 3.

### Immunoprecipitation

Immunoprecipitation was performed as previously described (Lim et al., 2015). Briefly, cleared cell lysates were incubated with 1µg WDR62, CEP170 or c-Myc antibodies conjugated to Protein-A agarose beads overnight at 4°C on an end-to-end rotator. After washing thoroughly with RIPA buffer, bound proteins were eluted with Laemelli buffer and separated by SDS-PAGE before immunoblotting.

### Live cell imaging

AD293 cells were seeded at 2×10^5^ cells per wells on a 35mm imaging dish (Ibidi µ-Dish 35 mm, high, 81156) in phenol red-free DMEM media (Gibco) supplemented with 10% FBS (Hyclone) and 1% penicillin-streptomycin (Gibco) to mitigate background fluorescence during imaging. Live cell imaging was performed on the Leica SP8 DMi8 inverted confocal microscope equipped with a Leica HC PL APO 20x 0.75 objective (Leica Microsystems). Temperature was maintained at 37°C and 5% CO2 in an enclosed incubation chamber. Specific areas of interest were selected using the Leica LAS X software (RRID:SCR_013673) and images were captured at 4 minutes interval for 14 hours. Analysis of mitotic duration was performed using the Leica LAS X software version 3.7.6.25997 (RRID:SCR_013673), Microsoft Excel and graphed using Graphpad Prism version 9.3 (RRID:SCR_002798).

### Statistics and Reproducibility

No sample size calculation was taken to predetermine sample size. All attempts at replication were successful, therefore no data was excluded for analysis. The biological *n* number for each experiment is detailed in the figure legends. All data were graphed as mean ± S.E.M. and statistically analysed (two-tailed unpaired student t-test or Two-way ANOVA) using Graphpad Prism version 9.3 (RRID:SCR_002798).

## RESULTS

### MAPKBP1 is localised to the subdistal appendages of the mother centriole

MAPKBP1 and WDR62 share structural similarities and can form heterodimers during stress response (Wasserman et al., 2010). They also share the same localisation at the centrosomes and basal body of the primary cilium (Jayaraman et al., 2016; Macia et al., 2017; Schonauer et al., 2020; Shohayeb et al., 2019). Therefore, we examined MAPKBP1 localisation in HeLa cells by expressing MAPKBP1 tagged at the N-terminus with GFP (GFP-MAPKBP1, **Figure 1A**) or by immunostaining endogenous MAPKBP1 using a commercially available MAPKBP1 antibody (**Figure 1B**) and showed localisation to centrosomes marked by γ-Tubulin. As a control, we expressed GFP-WDR62 in HeLa cells which was localised predominantly to the cytoplasm in interphase cells (**Figure 1A**) consistent with our prior findings (Bogoyevitch et al., 2012). We next confirmed that MAPKBP1 is located specifically on mother centrioles. We showed that GFP-MAPKBP1 co-localised with CEP170 or Ninein, known subdistal appendages proteins and mother centriole markers (**Figure 1C**, *upper and middle rows*). By contrast, GFP-MAPKBP1 did not co-localise with PCM1 which marks centriolar satellites (**Figure 1C**, *lower row*). Similarly, GFP-MAPKBP1 co-stained with γ-Tubulin at the basal body of the primary cilium (axoneme marked by ARL13B) (**Figure 1D**) consistent with localisation to the mother centriole. We have also examined GFP-MAPKBP1 localisation during prometaphase and metaphase, which showed MAPKBP1 localisation to the spindle poles (**Figure 1E**). This observation was confirmed using the commercial MAPKBP1 antibody, which showed similar localisation as WDR62 at the centrosomes during late S-phase and at the spindle pole during metaphase (**Supplementary Figure 1**), consistent with the literature (Macia et al., 2017; Schonauer et al., 2020).

**Figure 1.**
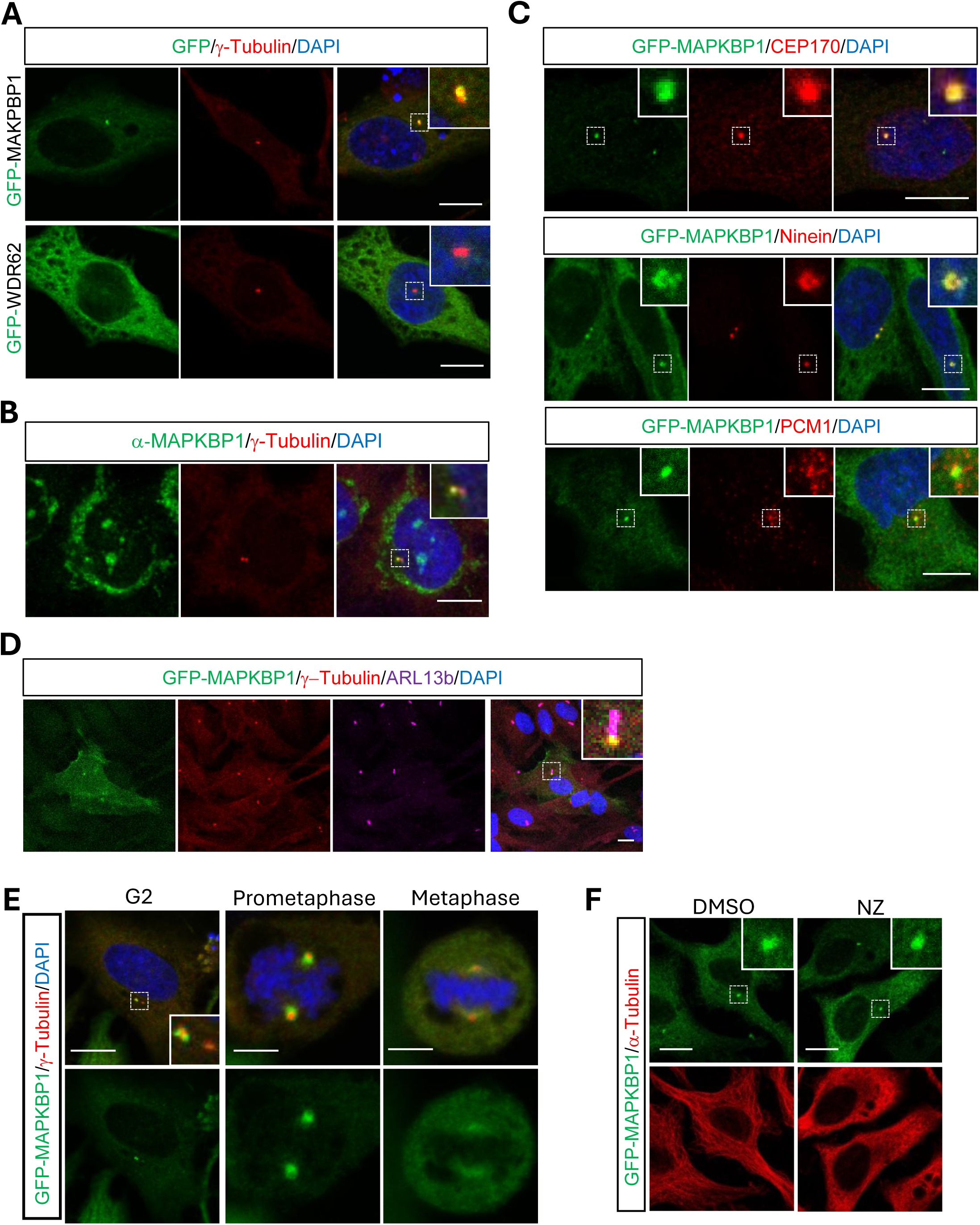
MAPKBP1 is localised to the mother centriole in interphase and to spindle poles in mitosis. **A.** Representative images of GFP tagged MAPKBP1 (GFP-MAPKBP1, green) co-localises with the centrosomes (marked by γ-Tubulin, red) in AD293 cells. GFP-tagged WDR62 (green) was used as a positive control for transfection. **B.** MAPKBP1 (using MAPKBP1 antibody, green) co-localises with the centrosomes (marked by γ-Tubulin, red). **C.** GFP-MAPKBP1 co-localises with known mother centriole markers – CEP170, Ninein, and PCM marker - PCM1 (all red). **D.** GFP-MAPKBP1 (green) co-localises with the basal body of the primary cilia, which is the centrosome (marked by γ-Tubulin, red). The axoneme of the primary cilia is marked by ARL13B (magenta). E. GFP-MAPKBP1 (green) is localised to the spindle poles (marked by γ-Tubulin, red) during prometaphase and metaphase. **F.** AD293 cells were treated with Nocodazole (NZ, 10 μM) or control DMSO for 30 mins to depolymerise microtubules (marked by α-Tubulin, red) and GFP-MAPKBP1 (green) localisation is unaffected. All cells were counterstained with DAPI (blue). Scale bars = 5µm.

To examine if MAPKBP1 centrosomal localisation is microtubule dependent, we treated HeLa cells with nocodazole, a small molecule that causes rapid depolymerisation of the microtubules by binding to tubulin and preventing microtubule polymerisation (Jordan et al., 1992). Nocodazole (**NZ**) treatment ablated α-tubulin-stained microtubules but did not affect MAPKBP1 localisation to the centrosome (**Figure 1F**). This observation is further consistent with MAPKBP1 localisation to centrioles and not the PCM or microtubule filaments. Taken together, our findings support previous research that MAPKBP1 is a centrosomal protein localised specifically to the subdistal appendages of the mother centriole (Macia et al., 2017; Schonauer et al., 2020).

### WDR62 is required for CEP170-MAPKBP1 interaction

While MAPKBP1 co-localises with CEP170 on mother centrioles and WDR62-CEP170 interactions is implicated in primary cilia regulation (Zhang et al., 2019), the interaction between MAPKBP1 and CEP170 interaction has not been previously characterised. Thus, we investigated MAPKBP1, WDR62 and CEP170 interactions in AD293 cells by co-immunoprecipitation (co-IP) studies. Through immunoprecipitation with WDR62 antibody, we showed that WDR62 was associated with MAPKBP1 and CEP170 (**Figure 2A**). As controls, we tested un-related centrosomal proteins, such as ch-TOG and CEP63, and showed that these proteins did not immunoprecipitate with WDR62 (**Figure 2A**). We next performed co-IP with CEP170 antibody showed that CEP170 immunoprecipitated WDR62 and MAPKBP1 (**Figure 2B**). To determine if WDR62 was required for CEP170 and MAPKBP to interact, we performed co-IP of WDR62 KO AD293 cell lysates (WDR62 KO) with CEP170 antibodies (**Figure 2C** and **Supplementary Figure 2A**). In the absence of WDR62 expression, we showed a loss of MAPKBP1 co-IP with CEP170 (**Figure 2C** and **Supplementary Figure 2A**). By contrast, the depletion of MAKPBP1 using siRNA (*siMAPKBP1*) did not alter the co-immunoprecipitation of WDR62 with CEP170 (**Figure 2D** and **Supplementary Figure 2B**). Likewise, depletion of CEP170 with siRNA (*siCEP170*) did not affect the interaction between WDR62 and MAPKBP1 (**Figure 2E** and **Supplementary Figure 2D**). Finally, we tested whether WDR62 missense mutations identified in primary microcephaly patients, WDR62 V65M and R438H, altered the interaction with CEP170 and MAPKBP1. We ectopically expressed Myc-tagged wild-type WDR62 (WDR62 WT) and showed that Myc pulldown, co-precipitated CEP170 and MAPKBP1. In addition, WDR62 V65M or R438H mutations did not impact the co-precipitation of MAPKBP1 with WDR62 (**Figure 2F**, *right panel middle blot* and **Supplementary Figure 2E**). This is consistent with the C-terminal dimerization domain of WDR62 required for MAPKBP1 interaction (Cohen-Katsenelson et al., 2011). By contrast, CEP170 interaction with Myc-tagged WDR62 V65M or R438H mutants was reduced substantially compared to WDR62 WT (**Figure 2F**, *right panel upper blot*). This indicates that disease-causing WDR62 mutations specifically alter interaction with CEP170 which suggests WDR62 binding of CEP170 may be involved in neurodevelopment. Our findings indicate that WDR62-CEP170-MAPKBP1 forms a complex and MAPKBP1 binding of CEP170 requires WDR62.

**Figure 2.**
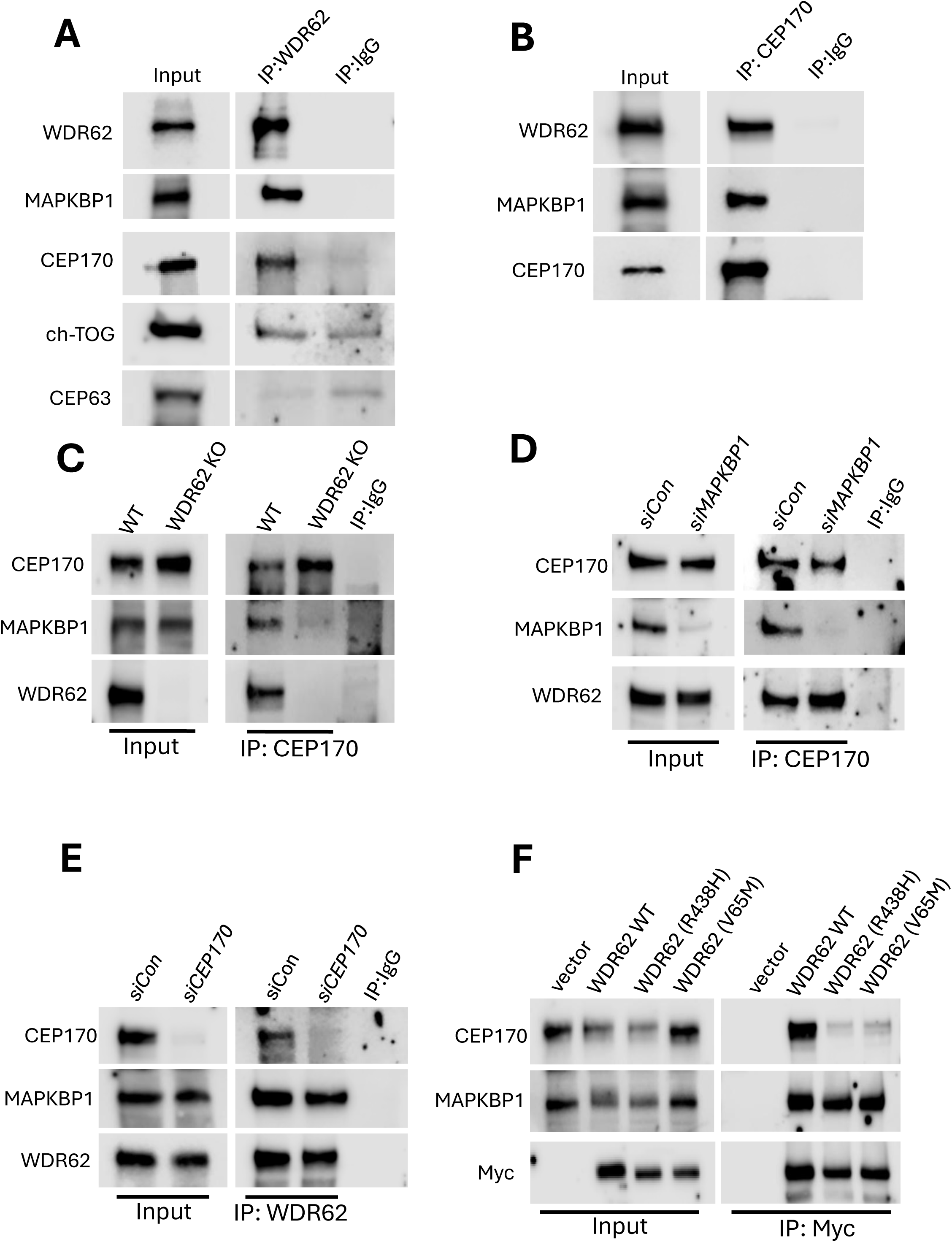
WDR62 interacts with MAPKBP1 and is required for MAPKBP1 to form a complex with CEP170. **A.** Immunoprecipitation using WDR62 antibody showing WDR62 interacts with MAPKBP1, CEP170 and did not interact with other centrosomal proteins, ch-TOG and CEP63, in asynchronous AD293 cells. **B.** Immunoprecipitation using CEP170 antibody showing CEP170 interacts with WDR62 and MAPKBP1. **C.** Immunoprecipitation using CEP170 antibody showing WDR62 depletion in WDR62 KO AD293 cells interfere with CEP170 and MAPKBP1 interaction. **D.** Immunoprecipitation using CEP170 antibody (IP: CEP170) showing MAPKBP1 knocked down (*siMAPKBP1*) did not interfere with CEP170 and WDR62 interaction. **E.** Immunoprecipitation using WDR62 antibody (IP: WDR62) showing CEP170 knocked down (*siCEP170*) did not affect WDR62 and MAPKBP1 interaction. **F.** Immunoprecipitation using c-Myc antibody (IP: Myc) showing Myc-tagged WDR62 carrying microcephaly associated mutations (V65M and R438H) affect WDR62-CEP170 interaction but not WDR62-MAPKBP1 interaction.

Next, we examined if siRNA silencing of CEP170, MAPKBP1 or WDR62 would affect the cellular localisation of this complex. We transfected AD293 cells with siRNAs against CEP170 (*siCEP170*), MAPKBP1 (*siMAPKBP1*) or WDR62 (*siWDR62*) and performed immunoblots to confirm siRNA mediated silencing of targeted protein expression (**Figure 3A** and **Supplementary Figure 3**). We also confirmed that *siCEP170* did not affect WDR62 and MAPKBP1 expression (**Figure 3A** and **Supplementary Figure 3**). Similarly, *siWDR62* did not lead to a decrease in CEP170 or MAPKBP1 compared to controls, and *siMAPKBP1* did not affect CEP170 protein levels but did lead to a modest decreased in WDR62 expression (**Figure 3A** and **Supplementary Figure 3**). We next evaluated the subcellular localisation of GFP-MAPKBP1 in *siCEP170* knockdown cells and showed reduced GFP-MAPKBP1 localisation to the centrosome of interphase cells and an absence of GFP-MAPKBP1 at the spindle poles of metaphase cells (**Figure 3B**). This indicated that CEP170 was required for MAPKBP1 localisation to centrosomes and spindle poles. By contrast, MAPKBP1 was not required for CEP170 localisation as *siMAPKBP1* knockdown did not change CEP170 localisation to the mother centriole in interphase or to mitotic spindles during metaphase (**Figure 3C**). Finally, we performed depletion of WDR62 in AD293 cell with *siWDR62*. While CEP170 localisation was unaltered with WDR62 depletion, the localisation of GFP-MAPKBP1 at the centrosome in interphase cells was lost when compared to *siCon*-treated cells (**Figure 3D**). Interestingly, the localisation of both GFP-MAPKBP1 and CEP170 to the spindle poles during metaphase were lost with WDR62 depletion (**Figure 3D**). Together with our co-IP experiments, these observations indicate that WDR62 is required for CEP170-MAPKBP1 interaction and localisation at the spindle poles.

**Figure 3.**
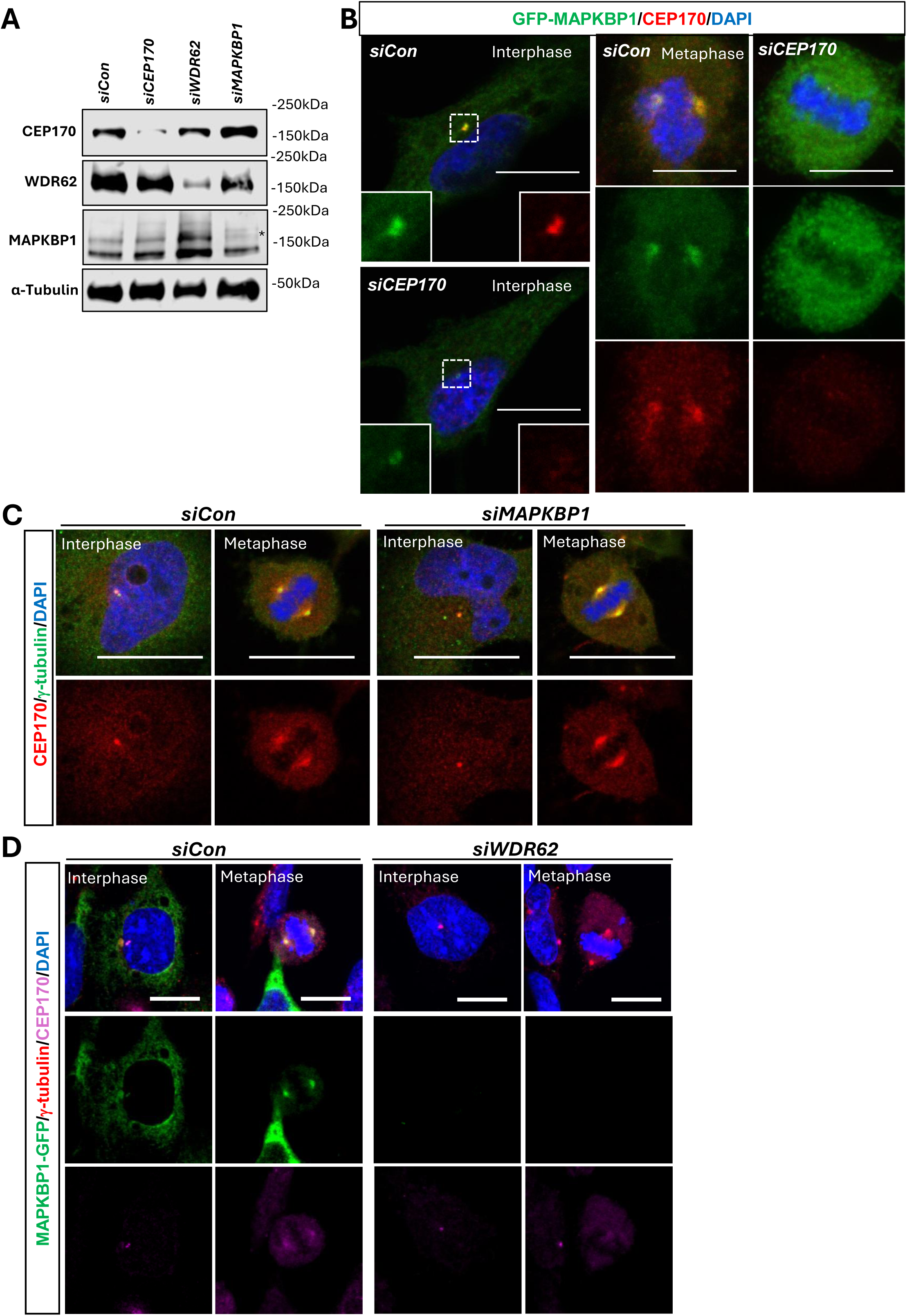
CEP170 and WDR62 is required for MAPKBP1 centrosome and spindle localisation. **A.** Representative CEP170, WDR62 and MAPKBP1 immunoblots confirming reduced targeted protein expression after siRNA treatment. CEP170, WDR62 and MAPKBP1 protein expression are unaffected with siRNA treatments (please see **Supplementary Figure 2** for raw data). **B.** GFP-MAPKBP1 (green) localisation is reduced at the centrosomes in interphase and metaphase when CEP170 (red) is silenced by *siCEP170*. **C.** CEP170 (red) localisation at the centrosome (marked by γ-Tubulin, green) is unaffected with *siMAPKBP1* treatment during interphase and metaphase. **D.** MAPKBP1-GFP (green) localisation at the centrosome (marked by γ-Tubulin, red) is lost with *siWDR62* treatment during interphase and metaphase. However, CEP170 (magenta) localisation is unaffected with *siWDR62* treatment. All cells were counterstained with DAPI (blue). Scale bars = 5μm.

### MAPKBP1 is not required for stress-induced JNK activation

WDR62 has been shown to interact with the activated form of JNK and recruit to stress granules during oxidative stress (Cohen-Katsenelson et al., 2011; Wasserman et al., 2010). Thus, we investigated if MAPKBP1 can interact with activated form of JNK in AD293/HeLa cells following stress treatment, namely hydrogen peroxide for oxidative stress and sorbitol for osmotic stress (Ding et al., 2021). Firstly, we overexpressed EGFP-MAPKBP1 or EGFP-WDR62 and HA-JNK1 or HA-JNK2 and showed that MAPKBP1 interacted less with JNK1 and JNK2 compared to WDR62 (**Figure 4A**). Next, we performed WDR62 immunoprecipitation following 1mM hydrogen peroxide treatment for 60min and demonstrated unchanged WDR62-MAPKBP1 interaction and protein levels (**Figure 4B**). We also performed WDR62 immunoprecipitation following 30min 0.5M sorbitol treatment and showed reduced WDR62 protein expression; however, WDR62-MAPKBP1 interaction remained unchanged (**Figure 4C**). In addition, we treated the cells with JNK inhibitor (JNK VIII) during hydrogen peroxide treatment and showed unchanged MAPKBP1, WDR62, total JNK (JNK 1 /2) and c-Jun protein levels (**Figure 4D**). Phosphorylation of JNK1/2 (p-JNK1/2) and c-Jun (p-c-Jun) were detected following stress treatment in DMSO-treated control as expected but were reduced or abolish in the presence of JNK inhibitor as expected (**Figure 4D**). We further investigated if silencing both MAPKBP1 and WDR62 would affect JNK phosphorylation during oxidative or osmotic stress, and our results showed that p-JNK1/2 and p-c-Jun were unaffected (**Figure 4E** and **4F**), suggesting that MAPKBP1 is not involved in stress-induced JNK and c-Jun phosphorylation.

**Figure 4.**
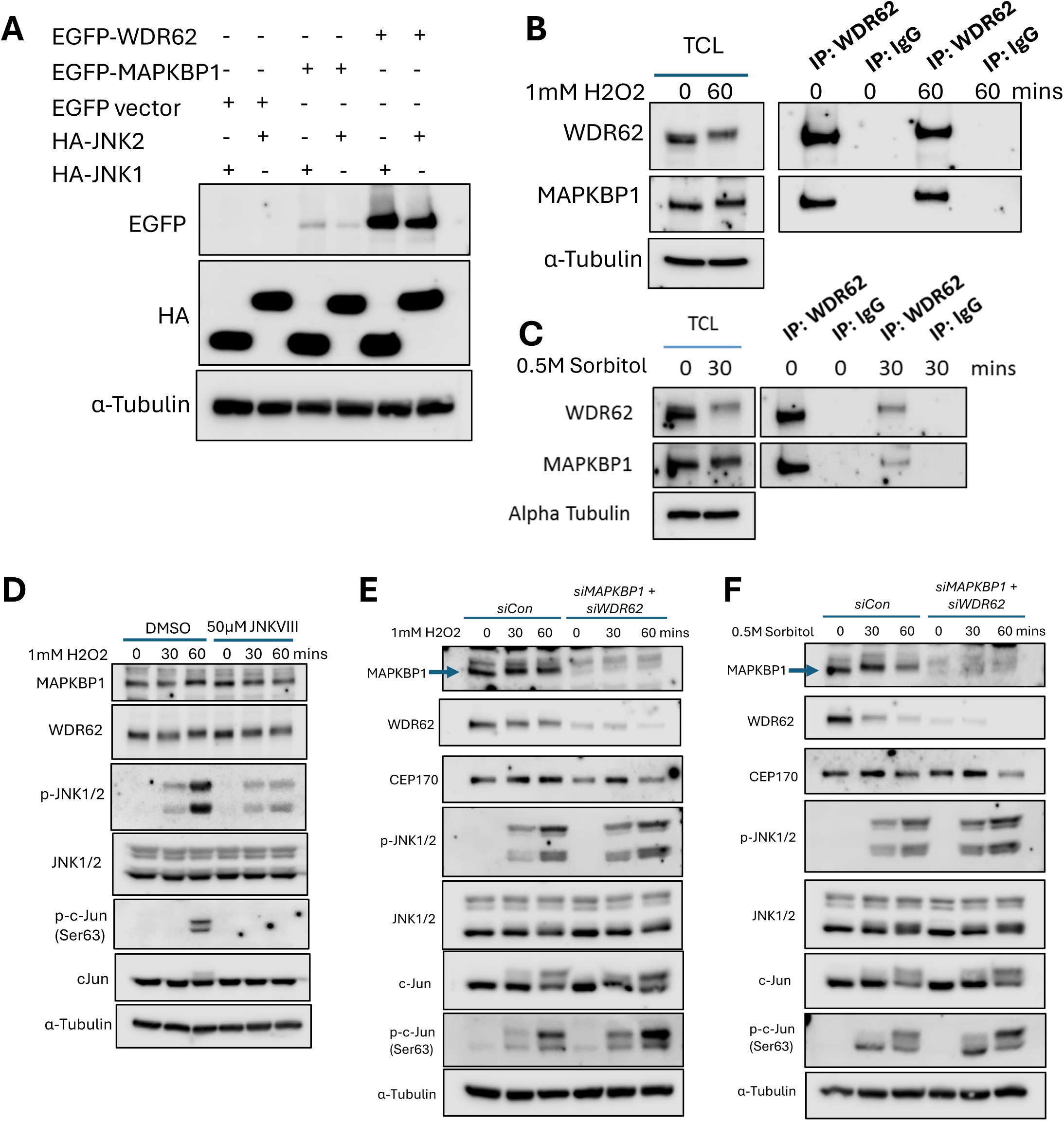
MAPKBP1 does not interact with JNK and is not required for stress-induced JNK activation. **A.** Immunoprecipitation of EGFP-WDR62, EGFP-MAPKBP1, HA-JNK1, HA-JNK2 and EGFP vector alone in AD293 cell lysates showing strong WDR62 association with JNK1 and JNK2, however, MAPKBP1 shows weak association with JNK1 and JNK2. **B.** AD293 cells were treated with 60 min 1mM H2O2 and immunoprecipitation shows unchanged WDR62 and MAPKBP1 interaction during oxidative stress compared to untreated. **C.** AD293 cells were treated with 30 min 0.5M sorbitol and immunoprecipitation shows weak WDR62 and MAPKBP1 interaction during osmotic stress compared to untreated. **D.** AD293 cells were treated with 50μM JNK inhibitor (JNKVIII) during 30 min and 60 min 1mM H2O2 treatment and immunoblot shows unchanged MAPKBP1, WDR62, total JNK1/2 and total c-Jun protein expression. Reduced phospho-JNK1/2 (Thr183/Tyr185) and phospho-c-Jun (Ser63) in the presence of JNKVIII as expected. **E.** AD293 cells were treated with *siMAPKBP1* and *siWDR62* during 30 min and 60 min 1mM H2O2 treatment and immunoblot shows reduced MAPKBP1 and WDR62 as expected, and unchanged CEP170, phospho-JNK1/2 (Thr183/Tyr185), total JNK1/2, total c-Jun and phospho-c-Jun (Ser63). **F.** AD293 cells were treated with *siMAPKBP1* and *siWDR62* during 30 min and 60 min 0.5M sorbitol treatment and immunoblot shows reduced MAPKBP1 and WDR62 as expected, and unchanged CEP170, phospho-JNK1/2 (Thr183/Tyr185), total JNK1/2, total c-Jun and phospho-c-Jun (Ser63). α-Tubulin was used as loading control in A, D, E and F.

### MAPKBP1 and WDR62 loss causes severe mitotic spindle defects

WDR62 is involved in the regulation of the bipolar mitotic spindle with WDR62 loss or mutation causing spindle defects and delayed mitosis (Lim et al., 2016). To determine if MAPKBP1 is also similarly required for spindle organisation, we depleted MAPKBP1 with siRNA and evaluated mitotic spindle assembly by staining with Pericentrin and α-Tubulin to label centrosomes and spindle microtubules respectively (Dictenberg et al., 1998). We quantified the number of normal bipolar mitotic spindles and compared this with bipolar spindles with mild or severe defects (*representative images* in **Figure 5A**). We found that MAPKBP1 depletion (*siMAPKBP1*) resulted in spindle defects comparable to WDR62 KO. Namely, in *siMAPKBP1* cells, we counted 26% of mitotic cells had mild or severe spindle defects compared to 41% in WDR62 KO cells (**Figure 5B**). By comparison, we did not detect abnormal spindles in AD293 cells transfected with control siRNA (*siCon*, **Figure 5B**). These findings indicate that MAPKBP1 has spindle regulatory functions similar to WDR62. To determine if MAPKBP1 loss would further exacerbate spindle defects in the absence of WDR62, we utilised siRNA to deplete MAPKBP1 in WDR62 KO cells (*siMAPKBP1*+WDR62 KO). This manipulation resulted in a substantial increase in spindle abnormalities with most cells (89%) exhibiting mild to several spindle defects with co-depletion of both MAPKBP1 and WDR62 (**Figure 5B**). This finding suggests a degree of functional redundancy between MAPKBP1 and WDR62 in mitotic and bipolar spindle regulation.

**Figure 5.**
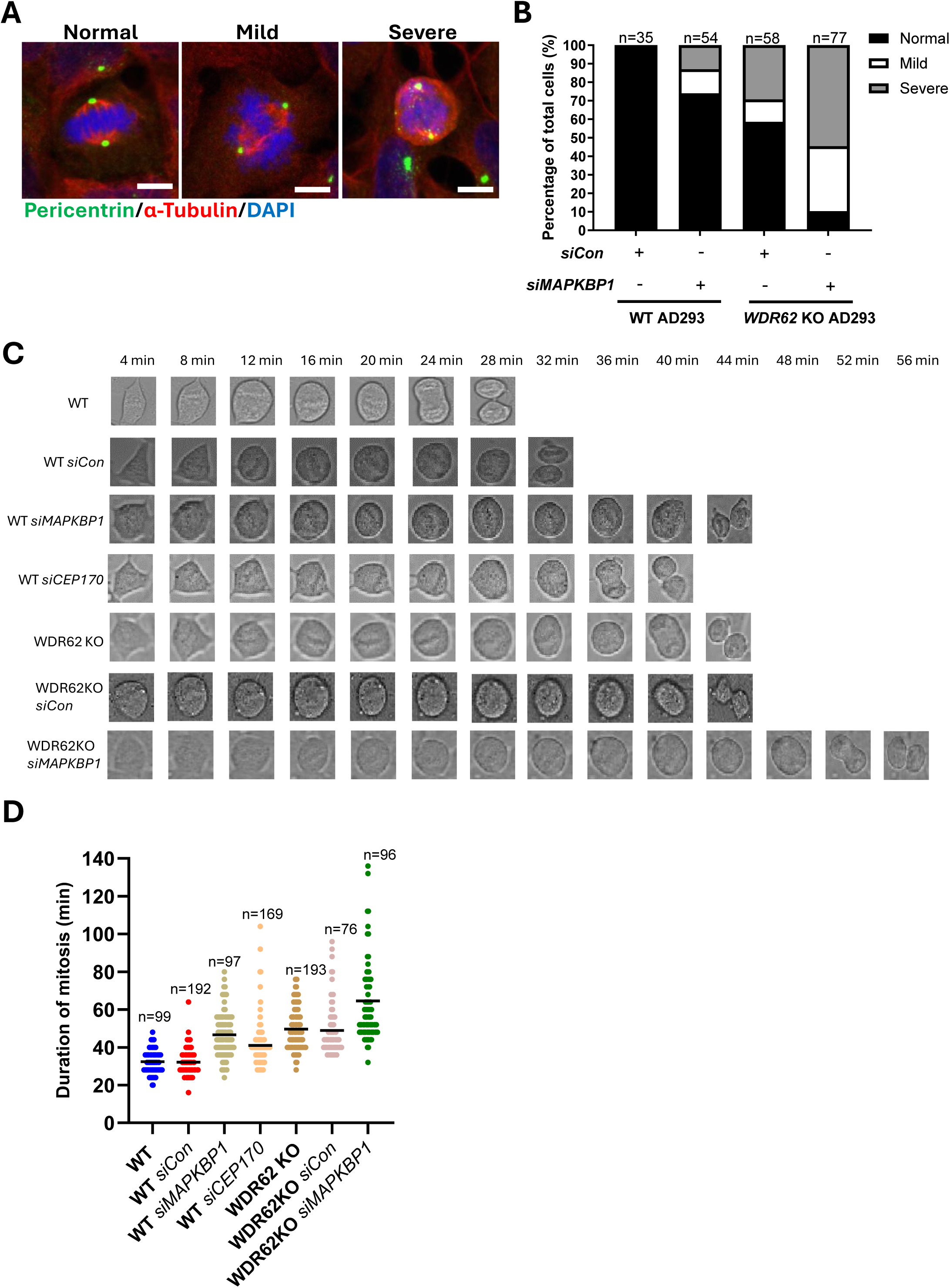
MAPKBP1 and WDR62 have overlapping functions and are partially redundant in spindle and mitotic regulation. **A.** Representative images of Pericentrin (green), α-Tubulin (red) co-staining illustrating normal, mild and severe defective mitotic spindles radiating from the bipolar spindle poles at metaphase. Cells were counterstained with DAPI (blue). Scale bar = 5 μm. **B.** Quantification of normal, mild and severe defective mitotic spindles from A in *siCon* WT AD293, *siMAPKBP1* WT AD293, *siCon*-WDR62 KO AD293, and *siMAPKBP1*-WDR62 KO AD293 cells. The total number of cells analysed are noted above the bar. **C.** Representative still images taken every 4 mins of untransfected (WT), *siCon*, *siMAPKBP1*, *siCEP170*, WDR62 KO, *siCon*-WDR62 KO and *siMAPKBP1*-WDR62 KO dividing cells. **D.** Quantification of mitosis duration (mins) from C. The total number of cells analysed at noted above the data. **E.** Quantification of the percentage of cells showing blebbing. The total number of cells analysed are noted above the bar. All data represent mean ± SEM.

Furthermore, we examined the impact of MAPKBP1 loss on mitotic progression. We performed live cell brightfield imaging to determine mitotic duration measured from when cells round up, as they enter mitosis, to when cells bifurcate and exit telophase (**Figure 5C**). Untreated AD293 cells or cells transfected with scrambled siRNA (*siCon*) completed mitosis with an average duration of 30 mins (**Figure 5C** and **5D**). In agreement with our previous findings, cells without WDR62 (WDR62 KO) took slightly longer (44 mins) to complete mitosis (**Figure 5C** and **5D**). We also found that siRNA depletion of CEP170 (*siCEP170*) or MAPKBP1 (*siMAPKBP1*) resulted in similar delay in mitotic progression (**Figure 5C** and **5D**), which reinforces a shared mechanism in mitotic regulation for these interacting partners. In findings reflecting the effects on spindle organisation, co-depletion of MAPKBP1 in the absence of WDR62 led to marked increase in mitotic duration (56 min) compared to loss of either WD-40 repeat protein alone (**Figure 5C** and **5D**). Collectively, these data suggest that the severe mitotic spindle defects are associated with a delay in mitosis.

### MAPKBP1 maintains pericentriolar matrix integrity

The centrosome’s intact structure and pericentriolar matrix (**PCM**) integrity provide a platform for microtubule nucleation essential for mitosis (Woodruff et al., 2017). Astral microtubules help to position and orient the spindle poles by linking the centrosomes to the cell cortex/membrane (Tame et al., 2014). Centrosome defects such as PCM fragmentation (structural integrity defect) and unseparated centrosomes (separation defect) during mitosis can sometimes be more obvious in the absence of dynamic microtubules (Hornick et al., 2008; Yang et al., 2009). Thus, we next investigated the effect of MAPKBP1 and WDR62 loss on centrosomes following inhibiting microtubule dynamics via taxol treatment. We transfected *siCon*, *siMAPKBP1* and/or *siCEP170* to WT and WDR62 KO AD293 cells and performed taxol treatment for 30mins. Then we analysed for centrosome and astral microtubules by immunostaining for pericentin and α-Tubulin respectively. Representative images of normal, fragmented centrosome and unseparated centrosome are illustrated in **Figure 6A**. In background of taxol-suppressed microtubule dynamics, we quantitated the percentage of cells with fragmented centrosomes and showed a significant increase in WDR62 KO cells (30%, *p*=0.0404) and *siMAPKBP1*-WDR62KO cells (36%, *p*=0.0079) compared to *siCon* (17%) (**Figure 6B**). There was also a significant increase in the percentage of WDR62 KO cells with unseparated centrosomes (36%, *p*=0.0265) compared to *siCon* (8%) (**Figure 6C**). *siMAPKBP1* depletion led to a subtle centrosome fragmentation effect (**Figure 6B**), but the number of unseparated centrosomes in mitosis was not significant elevated with MAPKBP1 depletion (**Figure 6C**). Given MAPKBP1 loss led specifically to fragmented centrosomes, our findings indicate that MAPKBP1 is involved in maintaining PCM cohesion and integrity.

**Figure 6.**
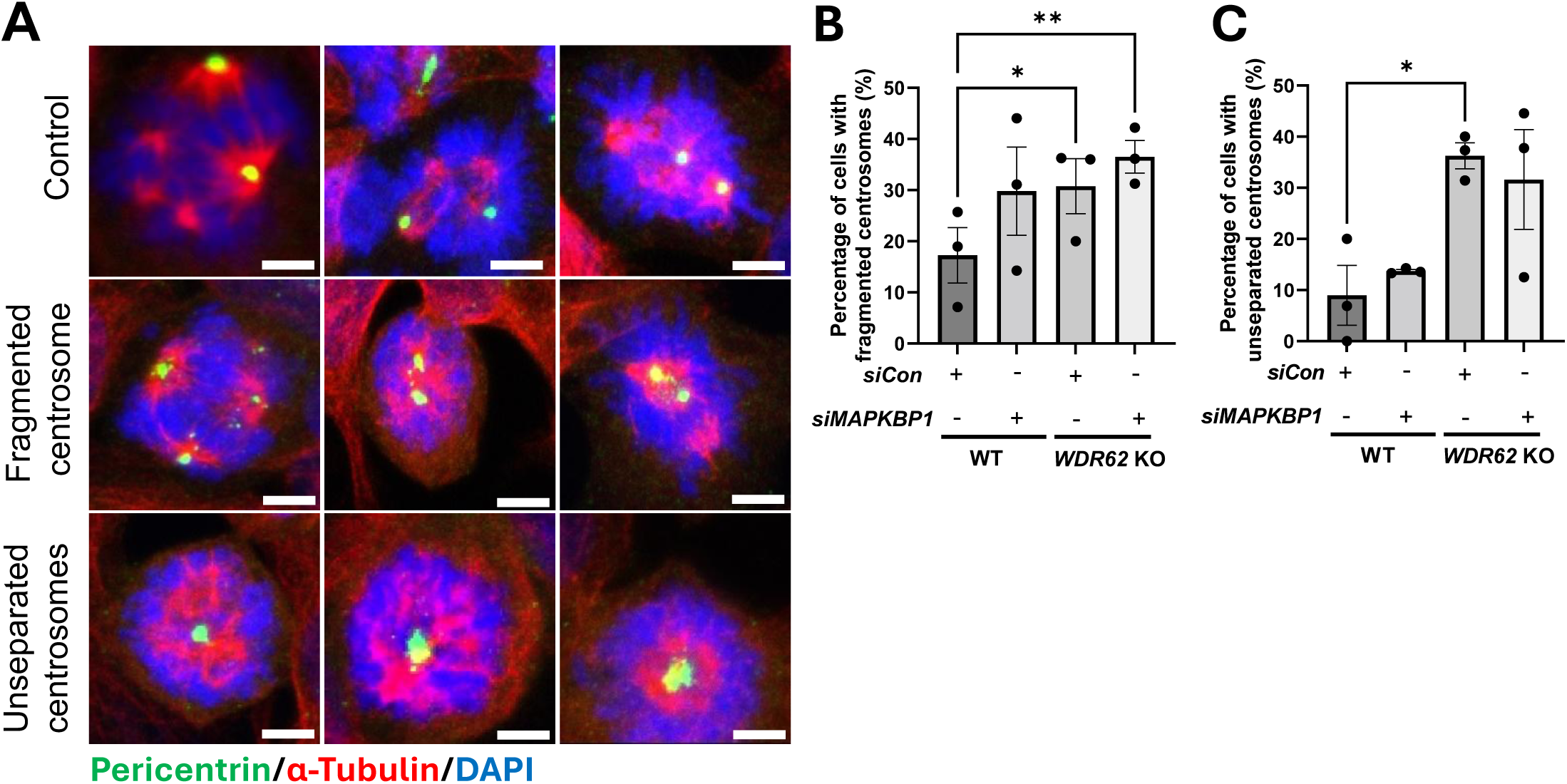
MAPKBP1 and WDR62 have parallel converging functions in centrosome cohesion. **A.** Representative images of Pericentrin (green), α-Tubulin (red) co-immunofluorescence illustrating normal, fragmented centrosome, and unseparated centrosome. Cells were counterstained with DAPI (blue). Scale bar = 5 µm. **B&C.** Percentage of cells showing fragmented centrosomes (**B**) and unseparated centrosomes (**C**) in *siCon* WT, *siMAPKBP1* WT, *siCon*-WDR62 KO, *siMAPKBP1*-WDR62 KO AD293 cells. A total of 143 *siCon* WT, 118 *siMAPKBP1* WT, 176 *siCon*-WDR62 KO, 144 *siMAPKBP1*-WDR62 KO AD293 cells counted from 3 independent experiments. All data represent mean ± SEM.

## DISCUSSION

MAPKBP1 has previously been shown to associate with centrosomes, mitotic spindles and the basal body of primary cilium (Macia et al., 2017; Schonauer et al., 2020). In this study, we have further characterised MAPKBP1 localisation at the subdistal appendages of the mother centrioles, and in the pericentriolar matrix (**PCM**) of the spindle poles during metaphase (**Figures 1E, 3B, 3D** and **Supplementary Figure 1**). MAPKBP1 shares the strongest overlap with WDR62 during metaphase at the spindle poles (**Supplementary Figure 1**) and with CEP170 at the mother centriole in during interphase (**Figure 3B** and **3D**). It is also recruited to the centrosomes by WDR62 and CEP170 as depletion of either protein has led to reduced MAPKBP1 recruitment (**Figure 3B** and **3D**). The subdistal appendages are mainly responsible for anchoring the minus end of microtubules to the centrosomes to form the microtubule organising centre, mitotic spindle regulation and maintenance of ciliary signalling (reviewed by (Ma et al., 2023)). During prometaphase and metaphase, subdistal appendages proteins such as CEP170 and Ninein have been reported to translocate to the PCM for spatial positioning of the spindle poles (Chen et al., 2003; Guarguaglini et al., 2005). Unlike WDR62, MAPKBP1 does not appear to associate with the microtubules, as inhibition of microtubule dynamics did not affect MAPKBP1 centrosomal localisation (**Figure 1F**). Therefore, based on intracellular localisation, MAPKBP1 is highly likely a subdistal appendage protein.

We have also interrogated MAPKBP1 function in this study. This is the first time that WDR62, MAPKBP1 and CEP170 have been shown to exist as a complex, where CEP170-MAPKBP1 interaction is mediated by WDR62 (**Figure 2C**). WDR62 has been studied extensively in regulating mitotic spindle and astral microtubule formation and dynamics, and also in spindle orientation (Bogoyevitch et al., 2012; Guerreiro et al., 2021; Huang et al., 2021; Lim et al., 2015; Miyamoto et al., 2017; Qin et al., 2019). Furthermore, spindle/astral microtubule disorganisation has been shown to cause spindle misorientation, activate spindle assembly checkpoint and delayed mitosis, as well as increased membrane blebbing in mitotic cells (Bogoyevitch et al., 2012; Chen et al., 2014; Kiyomitsu & Cheeseman, 2013; Rankin & Wordeman, 2010; Tame et al., 2014). Interestingly, MAPKBP1 depletion showed mitotic spindle defects and delayed mitosis similar to WDR62 KO (**Figure 5A** to **5D**) (Bogoyevitch et al., 2012; Farag et al., 2013; Lim et al., 2015; Qin et al., 2019). There also appears to be some additive effects in spindle defects and prolonged duration of mitosis when both MAPKBP1 and WDR62 are depleted (**Figure 5B** and **5D**). These observations suggest that MAPKBP1 and WDR62 may share overlapping function and/or heterodimerise at the spindle pole to stabilise mitotic spindles and astral microtubules during metaphase. Future experiments may investigate MAPKBP1 and WDR62 heterodimerisation at the mitotic spindles using a structural biology approach.

WDR62 and MAPKBP1 also diverge in their cellular functions. WDR62 is required for efficient stress-induced JNK activation (Wasserman et al., 2010), whereas MAPKBP1 is not, as shown in this study. Stress-induced JNK activation is mediated by ASK1, MEKK1, MLK3, and/or TAK1, which are typically cytoplasmic kinases (Brancho et al., 2005; Ichijo et al., 1997; Wang et al., 2001; Yujiri et al., 1998). Interestingly, MAPKBP1 is the only known subdistal appendage protein to associate with stress granules during cellular stress (Wasserman et al., 2010), despite subdistal appendage proteins generally functioning in centrosome structure and integrity rather than signal transduction (Chong et al., 2020; Ma et al., 2023). The mechanism by which MAPKBP1 contributes to stress granule formation or dynamics remains unclear and warrants further investigation. By contrast to its behaviour during stress, MAPKBP1 shares with WDR62 a C-terminal JNK-binding site (Cohen-Katsenelson et al., 2011), suggesting that both proteins can be phosphorylated by activated JNK at the centrosome or spindle poles. Activated JNK localises to centrosomes and phosphorylates WDR62 to regulate centrosome positioning and mitotic spindle formation (Bogoyevitch et al., 2012; Lim et al., 2015). A similar mechanism likely applies to MAPKBP1, as JNK-mediated phosphorylation has been shown to drive MAPKBP1 dissociation from the basal body of the primary cilium, promoting cilia disassembly (Findeisen et al., 2025). This suggests that MAPKBP1’s JNK-dependent regulation is context specific and distinct from its stress granule-associated role.

MAPKBP1 depletion mostly affected PCM cohesion but not centrosome separation (**Figure 6A** and **6B**). PCM components such as Pericentrin and γ-Tubulin have been shown to interact and form a lattice that helps to nucleate the minus end of microtubules (Dictenberg et al., 1998). In addition, depletion of Ninein, a well-known subdistal appendages protein, has been shown to affect the γ-Tubulin ring complex and subsequently uncoupled microtubule nucleation and anchorage at the centrosomes (Delgehyr et al., 2005). Furthermore, disruption of CEP128 at the subdistal appendages affects the spatial distribution of γ-Tubulin and loss of microtubule anchoring at the subdistal appendages (Chong et al., 2020). Therefore, some subdistal appendage proteins can also contribute to the organisation of PCM components. Since depleting MAPKBP1 affects PCM cohesion, it would be interesting to examine if MAPKBP1 provides a scaffold between subdistal appendages and the PCM components such as Pericentrin and/or γ-Tubulin. Taken together, MAPKBP1 depletion affects Pericentrin organisation at the PCM and subsequently microtubule nucleation for mitotic spindle formation and stability. Future studies should investigate the relationships between MAPKBP1 and other subdistal appendages and PCM proteins in regulating PCM cohesion and integrity.

In summary, MAPKBP1 is a subdistal appendages protein essential for PCM cohesion and subsequently microtubule nucleation for proper mitotic spindle formation. We have shown that CEP170 and WDR62 recruit MAPKBP1 in this process and WDR62 is required for CEP170-MAPKBP1 interaction. This study has emphasised how microtubules nucleation at the centrosomes are tightly regulated by subdistal appendages and scaffolding proteins to ensure proper chromosome alignment and synchronous chromosome segregation during cell division.

## AUTHOR CONTRIBUTION

**Uda HO:** Conceptualisation, methodology, investigation, formal analysis, visualisation, writing – original draft, writing – review and editing, supervision. **Yvonne YEAP:** Methodology, investigation, validation, formal analysis, visualisation. **Cheng-Hui GOH:** Investigation, validation, formal analysis, visualisation. **Peter NOAKES**: writing – reviewing and editing, resources. **Dominic NG:** Conceptualisation, project administration, writing – reviewing and editing, supervision, resources, funding acquisition.

## ACKNOWLEDGEMENT

This project was funded by National Health and Medical Research Council Australia (GNT1162652) awarded to A/Prof. Dominic Ng. Dr. Uda Ho was also supported by FightMND discovery grant (DIS: 202403-01216) awarded to A/Prof. Peter Noakes. We would like to thank the UQ-SBMS microscopy and analytical facility for their assistance.

## DATA AVAILABILITY

All data are included in this publication. Raw imaging files can be found in UQ-RDM/UQ eSpace (DOI link will be provided at final submission).

