## Supplementary Figures and Tables for "WDR62 and CEP170 recruit MAPKBP1 for pericentriolar material cohesion and mitotic spindle formation"

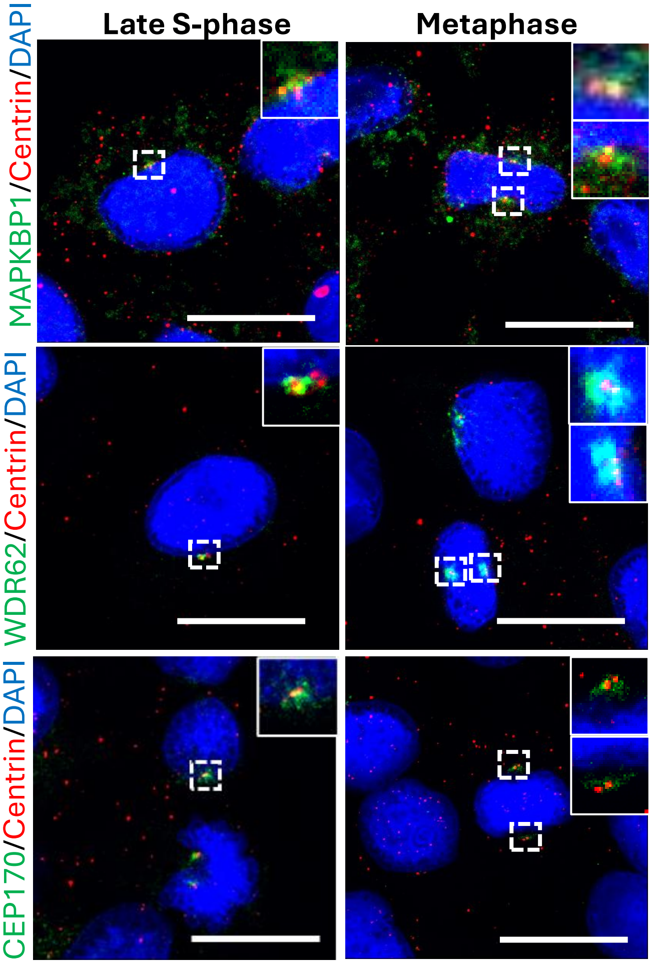

**Supplementary Figure 1. MAPKBP1, WDR62 and CEP170 localisation in HeLa cells.**
Representative images of MAPKBP1 (top panel, green), WDR62 (middle panel, green) and CEP170 (bottom panel, green) co-staining with the centriole marker, Centrin (red), in late S-phase and metaphase HeLa cells. All three proteins are localised to the centrosomes during late S-phase, whereas MAPKBP1 and WDR62 are also localised to the spindle poles during metaphase and CEP170 remains associated to the centrioles. Cells are counterstained with DAPI (blue). Scale bar = 20μm.

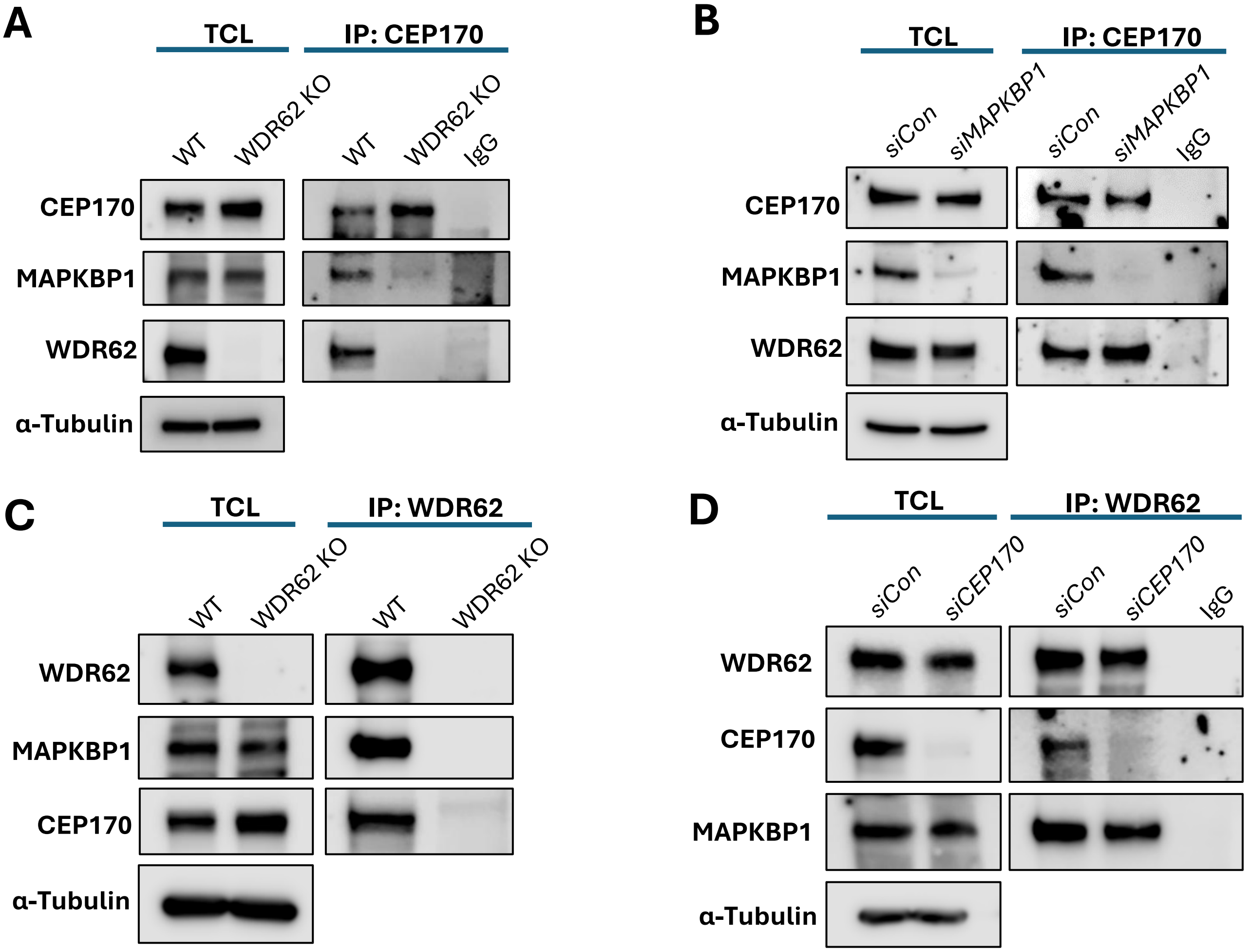

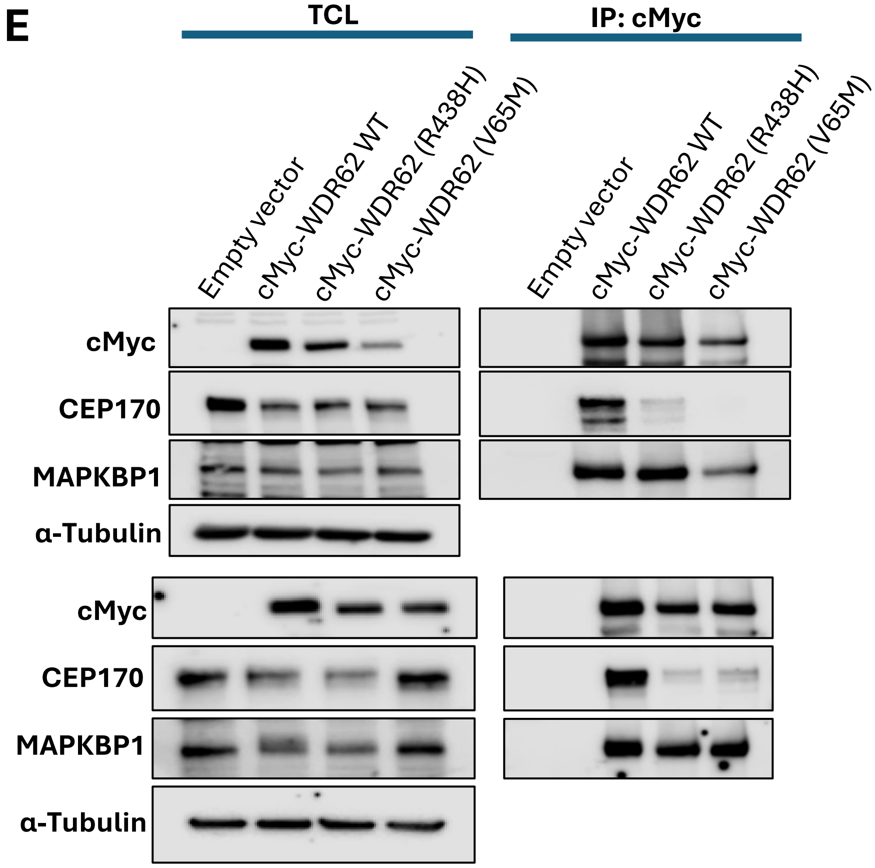

**Supplementary Figure 2. Immunoprecipitation data related to Figure 2.**

**A.** CEP170 immunoprecipitation (IP) in WDR62 KO AD293 cells showing no CEP170 association with MAPKBP1 in the absence of WDR62. Related to Figure 2C. **B.** CEP170 IP in siMAPKBP1-treated AD293 cells showing MAPKBP1 depletion did not affect CEP170-WDR62 interaction. Related to Figure 2D. **C.** WDR62 IP showing no CEP170-MAPKBP1 interaction in the absence of WDR62. **D.** WDR62 IP in siCEP170-treated AD293 cells showing depletion of CEP170 did not affect WDR62-MAPKBP1 interaction. Related to Figure 2E. **E.** cMyc IP showing reduced CEP170 association in WDR62 mutants (V65M, R438H) compared to wildtype, indicating CEP170 most likely associates with the N-terminus of WDR62. In contrast, WDR62 mutation did not affect WDR62-MAPKBP1 interaction most likely at the C-terminus. Related to Figure 2F.

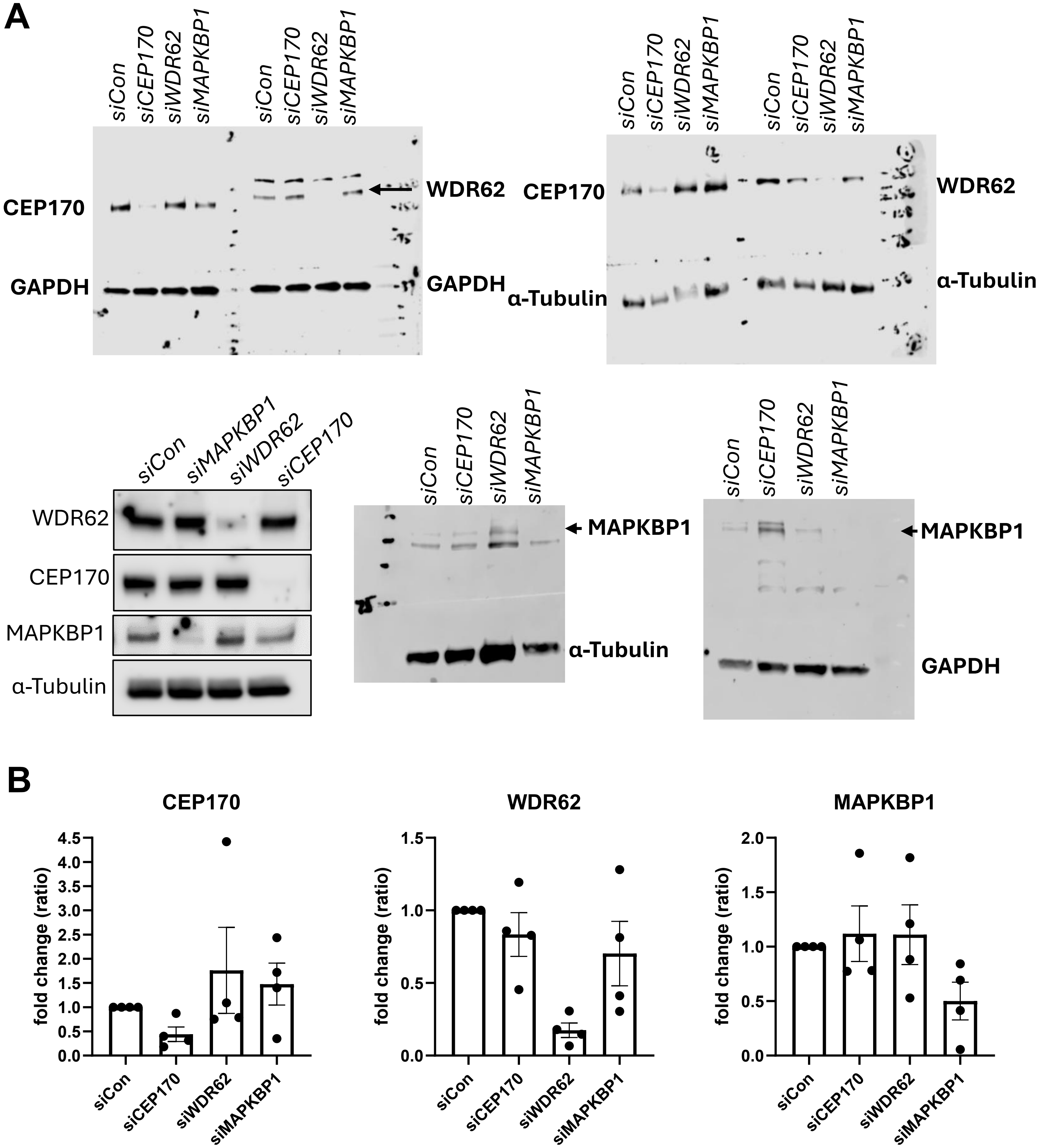

**Supplementary Figure 3. MAPKBP1, WDR62 and CEP170 western blots of siRNA experiments related to Figure 3.**

**A.** Raw blots of siRNA experiments in AD293 cells related to Figure 3A. **B.** Quantification of (A) confirmed siRNA mediated silencing of targeted protein expression normalised to loading controls (GAPDH or α-Tubulin). Data represents as mean ± SEM.

#### Supplementary Table 1. Plasmids and siRNA used in this paper.

| Name of siRNA or plasmid | Manufacturer and catalogue number | Final concentration used in experiment |
|---|---|---|
| Human MAPKBP1 siRNA | Dharmacon, L-022225-00-0005 | 20nM in AD293 cells |
| Human CEP170 siRNA | Dharmacon, L-021258-00-0005 | 20nM in AD293 cells, 60nM in HeLa cells |
| Human WDR62 siRNA | Dharmacon, J-031771-09-0020; J-031771-12-0020 | 20nM in HeLa cells |
| Scrambled siRNA | Dharmacon, D-001810-10-20 | 20nM in AD293 cells, 60nM in HeLa cells |
| pEGFP-N3 | Clontech, 6080-1 | 1µg per well in 6 well plate |
| pEGFP-MAPKBP1 | In house | 1µg per well in 6 well plate |
| pJX40-cmyc-WDR62-FL(WT) | In house (Lim et al., 2016) | 1µg per well in 6 well plate |
| pJX40-cmyc-WDR62-FL(V65M) | In house (Lim et al., 2016) | 1µg per well in 6 well plate |
| pJX40-cmyc-WDR62-FL(R438H) | In house (Lim et al., 2016) | 1µg per well in 6 well plate |

### Supplementary Table 2. Primary antibodies used in this paper.

| Name of antibody | Host species | Manufacturer, catalogue number and RRID | Dilution |
|---|---|---|---|
| **Primary antibodies** | | | |
| WDR62 | rabbit | Bethyl Laboratories, A301-560A (IF) (RRID:AB_1040044) A301-559A (WB, IP) (RRID:AB_1040043) | 1:200 Immunofluorescence (IF) 1:1000 Western blotting (WB) 1μg Immunoprecipitation (IP) |
| CEP170 | rabbit | Abcam, AB72505 (RRID:AB_1268101) | 1:200 IF 1:1000 WB 1μg IP |
| MAPKBP1 | rabbit | Proteintech, 17788-1-AP (RRID:AB_2235097) | 1:200 IF 1:1000 WB |
| Ninein | rabbit | Abcam, AB52473 (RRID:AB_881057) | 1:200 IF |
| Centrin | mouse | Millipore, 04-1624 (RRID:AB_10563501) | 1:400 IF |
| Pericentrin | rabbit | Abcam, AB4448 (RRID:AB_304461) | 1:200 IF |
| α-Tubulin | mouse | Sigma, T5168 (RRID:AB_477579) | 1:200 IF |
| γ-Tubulin | mouse | Sigma, T5326 (RRID:AB_532292) | 1:200 IF |
| PCM1 | rabbit | Cell Signaling, 5259S (RRID:AB_10556961) | 1:200 IF |
| ARL13B | rabbit | Proteintech, 17711-1-AP (RRID:AB_2060867) | 1:200 IF |
| CEP63 | rabbit | Millipore, 06-1292 (RRID:AB_10918481) | 1:1000 WB |
| Ch-TOG | rabbit | Abcam, AB86073 (RRID:AB_1924889) | 1:1000 WB |
| JNK1/2 | mouse | BD Pharmingen, 554285 (RRID:AB_395344) | 1:1000 WB |
| p-JNK1/2 (Thr183/Tyr185) | rabbit | Cell signaling, 612540 (RRID:AB_399837) | 1:1000 WB |
| c-Jun | rabbit | Cell signaling, 9165 (RRID:AB_2130165) | 1:1000 WB |
| p-c-Jun (Ser63) | rabbit | Cell signaling, 9261S (RRID:AB_2130162) | 1:1000 WB |
| GFP | chicken | Invitrogen, A10262 (RRID:AB_10262) | 1:500 IF |
| MYC | rabbit | Sigma, A7407 Santa Cruz Biotech, sc-789 (RRID:AB_631274) | 1:1000 WB 1μg IP |
| GAPDH | mouse | Abcam, AB9485 (RRID:AB_307275) | 1:1000 WB |
| IgG | rabbit | Santa Cruz, sc-2027 (RRID:AB_737197) | 1μg IP |

### Supplementary Table 3. Secondary antibodies used in this paper.

| Name of antibody | Host species | Manufacturer, catalogue number and RRID | Dilution |
|---|---|---|---|
| Mouse Alexa 488 | goat | Invitrogen, A11001 (RRID:AB_2534069) | 1:250 IF |
| Mouse Alexa 555 | goat | Invitrogen, A21424 (RRID:AB_141780) | 1:250 IF |
| Rabbit Alexa 488 | goat | Invitrogen, A11034 (RRID:AB_2576217) | 1:250 IF |
| Rabbit Alexa 555 | goat | Invitrogen, A21428 (RRID:AB_2535849) | 1:250 IF |
| Rabbit Alexa 647 | goat | Invitrogen, A21245 (RRID:AB_2535813) | 1:250 IF |
| Mouse HRP | goat | Pierce, PIE31430 | 1:5000 WB |
| Rabbit HRP | goat | Pierce, PIE31460 | 1:5000 WB |
| DAPI 5mg/ml | n/a | Sigma, D8417 | 1:10,000 IF |
